# Developmental genetic response of the zooplanktonic tunicate *Oikopleura dioica* to marine noise pollution

**DOI:** 10.64898/2026.05.28.728403

**Authors:** Eva R. Quintana, Nuria P. Torres-Águila, Ignasi Nou-Plana, Sissel Norland, Valentina Caorsi, Giorgio Blumer, Matteo Bozzo, Elettra Panarari, Giacomo Sabaddin, Simona Candiani, Irene Guarneri, Lucia Manni, Roberta Pennati, Filomena Ristoratore, Giovanni Zambon, Marios Chatzigeorgiou, Rosa Maria Alsina-Pagès, Cristian Cañestro

## Abstract

**Background:** Anthropogenic noise is an emerging threat to marine ecosystems, yet its effects on marine invertebrates, particularly zooplanktonic species, remain poorly understood. Despite increasing evidence of behavioral and physiological impacts in invertebrates, the effects of noise on embryonic development and the molecular mechanisms underlying acoustic responses remain largely unexplored. Here, to address this gap, we investigated the impact of high-intensity underwater noise exposure on embryogenesis of the appendicularian tunicate *Oikopleura dioica*, a cosmopolitan zooplanktonic tunicate that plays important ecological roles in marine trophic webs and carbon cycling. Under lab-controlled conditions, we examined the effects of experimental noise exposure on early embryogenesis at both morphological and transcriptomic levels using RNA-seq in 8-cell (8c) and early tailbud (ETB) stages.

**Results:** Noise exposure produced no significant increase in embryo malformations compared to non-exposed controls, indicating substantial phenotypic resilience under laboratory conditions. Interestingly, transcriptomic analyses revealed a rapid molecular response of 70 differentially expressed genes (DEG) already detectable after only 30 minutes of exposure at the 8-cell stage, which became markedly amplified with 700 DEGs by the ETB stage. Together, differential expression, GO enrichment, and co-expression network analyses identified coordinated regulation of processes associated with membrane homeostasis, pyrimidine/CTP metabolism, extracellular matrix organization, cytoskeletal architecture, RNA regulation, translational control, proteostasis, mitochondrial metabolism, and developmental pathways. Importantly, both developmental stages precede the formation of differentiated mechanosensory structures, suggesting that the observed responses are unlikely to reflect conventional sound perception.

**Conclusions:** These findings provide the first molecular characterization of noise effects during *O. dioica* embryogenesis and reveal an unexpected molecular sensitivity of *O. dioica* embryos to underwater noise despite preserved morphological development. The transcriptional signatures support a mechanobiological framework in which acoustic exposure may directly perturb cellular mechanical homeostasis through membrane-and cytoskeleton-associated processes, triggering compensatory stress-adaptation responses involving proteostasis, RNA regulation, and metabolic reprogramming. Together, these findings establish *O. dioica* as a valuable emerging model for investigating the developmental and evolutionary consequences of acoustic pollution in marine ecosystems.

## Background

Anthropogenic noise has become an increasingly important form of pollution, with detrimental effects on marine biodiversity and ocean health [1]. Growing evidence demonstrates that underwater noise impairs critical behaviours across marine animals, ranging from subtle communication disruption and avoidance responses [2–4], to pronounced physiological stress [5] and increased mortality rates [6]. For example, in goby fish (*Gobiusculus sp.* and *Pomatoschistus sp.*), courtship and successful spawning become impaired following noise exposure [7].

While most studies have traditionally focused on so-called “hearing species”, that is, animals such as marine fish and mammals with specialized auditory systems, invertebrate organisms remain largely overlooked despite possessing “sound-sensitive” structures that make them responsive to mechanical and acoustic cues [8]. Given the central ecological role of marine invertebrates, this limited understanding constrains our ability to predict how acoustic disturbance propagates through marine food webs [reviewed in 9]. Addressing this issue is critical for Ocean Health, as disruptions to marine communities can impair nutrient cycling and carbon sequestration, ultimately threatening food security and climate stability [10].

Evidence supporting that noise affect invertebrates include recent studies on copepods showing that exposure to intense acute noise increases mortality and induces developmental damage [11], while chronic exposure to boat noise impairs feeding and reproductive performance [12, 13]. Similarly, the amphipod *Corophium volutator* exhibits reduced bioturbation activity and shallower burial depths under low-frequency noise exposure [14]. In the sea urchin *Arbacia lixula*, acute high-intensity noise generated by seismic airguns triggers inflammation of the peristomial membrane and elicits a pronounced stress-related adaptive response [15]. Collectively, these behavioural and physiological effects represent a convergent manifestation stress-induced responses across marine invertebrates, in which acoustic disturbance alters energy allocation and organism-environment interactions.

In contrast to the impact of noise at the behavioural and physiological levels, our understanding of its impact at the genetic level remains poorly characterized and is largely limited to a few vertebrate species. Some recent studies show, for instance, that porpoises (*Neophocaena sp.*) upregulate the expression of immune genes and oxidative stress markers [16], while fish exhibit stress responses ranging from altered synaptic transmission genes in small yellow croaker (*Larimichthys polyactis*) [17] to changes in neuroplasticity-related genes in brook trout (*Salvelinus fontinalis*) [18]. During embryo development, early stages appear particularly vulnerable, with zebrafish exposed to high-intensity noise showing disruption of axon guidance and MAPK signaling pathways [19].

In invertebrates, although studies are still scarce, emerging evidence points to analogous molecular stress responses to those described in vertebrates. Horseshoe crabs (*Tachypleus tridentatus*), for instance, upregulate heat shock proteins and exhibit metabolic dysregulation in transcriptomic analyses [6], while silver-lipped pearl oysters (*Pinctada maxima*) exhibit long-lasting disruption of energy production pathways persisting months after seismic exposure (Dang et al., 2024). Sea cucumbers (*Apostichopus japonicus*) and mussels (*Mytilus spp.*) exhibit oxidative damage and immune suppression gene expression changes under continuous noise, respectively [20, 21]. These transcriptomic data suggest that behavioral and physiological disruptions observed in invertebrates reflect underlying molecular stress mechanisms similar to those describe in vertebrates, although detailed mechanistic understanding remains limited for most species.

Among transcriptomic studies investigating the impact of anthropogenic noise on marine invertebrates, zooplanktonic mesopelagic species remain largely underrepresented [22], leaving a significant gap in our understanding of the vulnerability of pelagic trophic webs to acoustic pollution. To fill this gap, we focus here on the cosmopolitan appendicularian *Oikopleura dioica*, one of the most abundant mesozooplanktonic organisms after copepods. *O. dioica* represents an ideal model for studying the effects of noise on zooplanktonic animals with short life cycles, from 5 to10 days depending on temperature, and a semelparous reproductive strategy, where mature individuals release eggs and sperm into the water column for external fertilization and die shortly thereafter. As a result, embryonic development occurs entirely in the environment and likely constitutes a particularly vulnerable stage to external perturbations such as marine noise pollution. This vulnerability is expected to be especially relevant in organisms with semelparous life strategies, as reproductive success depends on a single reproductive event and cannot be compensated by subsequent breeding cycles in different locations. Furthermore, *O. dioica* combines a simple and accessible morphology, rapid generation time, ease of laboratory culture, and experimental tractability [23], together with a high-quality, well-annotated genome that enables robust transcriptomic analyses [24, 25]. Ecologically, this filter-feeding tunicate plays a key role in marine biogeochemical cycles by producing marine snow through fecal pellets and discarded mucus houses, thereby influencing the vertical flux of organic matter and contributing to nutrient cycling and carbon sequestration [26]. Finally, its phylogenetic position at the base of the chordate lineage provides a valuable comparative framework for understanding how stress responses have evolved across vertebrates and invertebrates.

Here, we investigate the effects of anthropogenic noise on the embryonic development of *O. dioica,* at both morphological and genetic levels, contributing to narrowing the knowledge gap regarding the impacts of noise pollution on zooplanktonic organisms and marine ecosystems.

## Methods

### Laboratory culture of *Oikopleura dioica*

*O. dioica* specimens were acquired from animal lab colonies that have been maintained in our facility in the University of Barcelona for over five years. The founder individuals were originally obtained from the Mediterranean coast near Barcelona (Catalonia, Spain) and cultured as detailed in Martí-Solans, et. al. 2015 [27].

### Noise exposure and set-up experimental calibration of the tank

A glass tank aquarium of 130 cm long, 40 cm wide and 60 cm high was filled with 280 L of water, and an AKUASOUND_16 underwater speaker (MAUSOUND, San Damaso, Modena) was submerged located near one of the walls oriented towards the centre, and connected via a NX3000D power amplifier (Behringer, Willich, Germany) to a computer running Audacity software, from which the different noise profiles were reproduced (version 3.6, https://www.audacityteam.org/). *O. dioca* animals were placed in a 2 L glass beaker filled with 1,8 L of sea water suspended in the middle of the water column, hanging from a nylon rod from the ceiling of tank to avoid the transmission of vibration from the walls of the tank to the beaker (**Fig. 1**). The acoustic environment of the tank was characterized to ensure accurate and reproducible noise conditions by measuring sound pressure levels using an AS-1 hydrophone (Aquarian Audio, Anacortes, WA, USA). Specifically diving the tank into a grid of 3×4. Using this knowledge, we took measurements on the specific site where the animal was fixed (C2) in the tank while exposed to noise for the different playbacks, inside and outside the beaker and inside and outside the prototypes, assessing spatial variability in sound distribution and characterizing the acoustic signal propagation profile. The analysis demonstrated that, as expected, the sound pressure levels were not homogeneous within the tank (**Supplementary figure S1A**), as also reported by other studies, suggesting that choosing one central site in the tank, away from the wall is regarded as the best choice [28, 29]. Based on this, the position C2 (top and bottom, inside outside baker) was selected as the standardized exposure point for the present experiments (**Supplementary figure S1B**).

**Figure 1.**
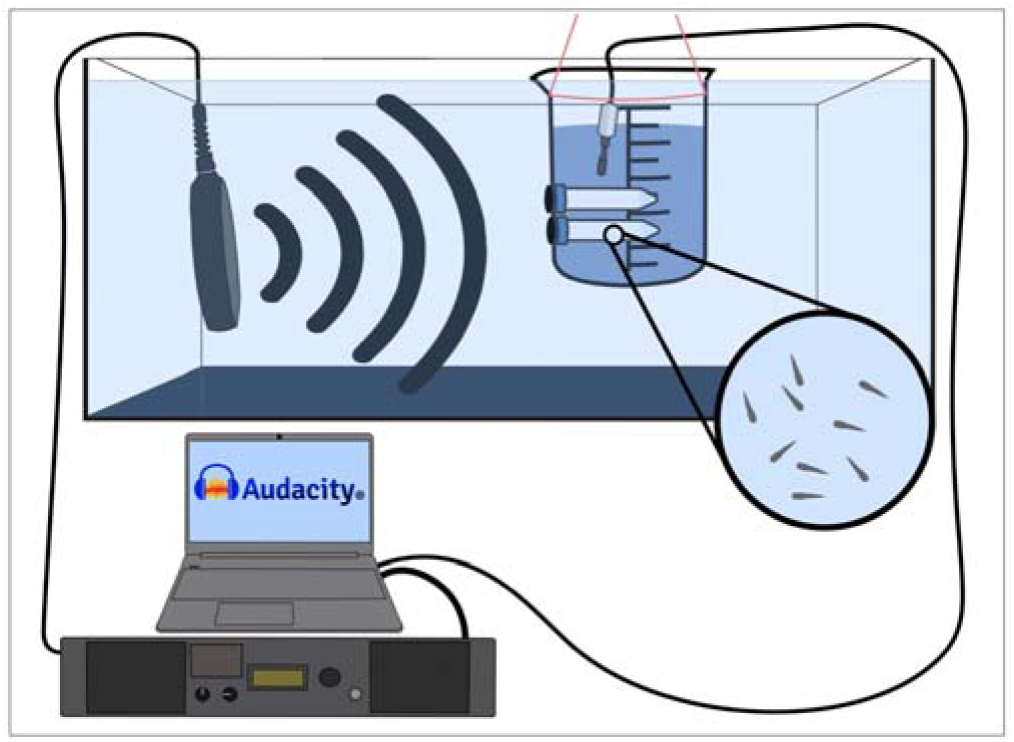
Outline of the set-up for sound exposure experiments to *O. dioica* embryos.

For noise exposure experiments, embryos were placed in a meshed chamber inside the beaker, which consisted of a windowed 15 mL falcon with 100 µm cell mesh allowing water and sound flow and placed horizontally regarding the speaker (**Fig. 1**).

Noise characterization was conducted for all types of noise files by using both pure tones (PT: 32.5 Hz, 125 Hz, 500 Hz and 1000 Hz) as well as noise bands (NB: 32.5-125 Hz, 125-500 Hz, 500-1000 Hz, 63-125 Hz), the latter being a proxy to real anthropogenic noise recommended by the Directorate-General for Environment of the European Commission (EC/NB:63-125). The noise PT signals and NB files were generated by La Salle – Universitat Ramon Llull using Audacity. Pure tones (PT) were created using the “Generate Tone” function. Noise bands (NB) were produced using the “Generate Noise” function (white noise) and subsequently filtered to obtain the desired frequency ranges. The noise of PT-and NB-files was played at the maximum intensity possible (153–163 dB re 1 µPa; 100 W RMS power with an impedance of 8 Ohms) aiming to investigate the most harmful possible impact of noise on the biology of *O. dioica*. All experimental exposures were subsequently conducted at a fixed position within the tank, corresponding to the calibrated exposure point. The control beaker was placed in the same room in which the experiment was being performed in order to share the same temperature and environmental conditions than treated animals, but far enough from the tank (>15 m away) to avoid the effect of sound or any potential surface vibration. In addition, background noise levels were recorded in the absence of experimental sound playback and consistently remained at least 20 dB re 1 µPa lower than the treatment levels throughout the recording period. These measurements validated the acoustic calibration of the experimental set-up and confirmed that experimental exposures were conducted under well-defined, stable, and controlled noise conditions.

### Phenotype and malformation rate after noise exposure during embryo development

For noise exposure of embryos, a pool of eggs from at least three females (i.e. 200-500 eggs) were fertilized with pooled sperm from at least three males as previously described in Martí-Solans, et. al. 2015 [27], to obtain genetically diverse embryos developing simultaneously and avoiding potential biologic bias towards one particular parent. Following visual confirmation of a good fertilization rate by the appearance of the polar body and synchronous first cleavage in most of the clutch, embryos were split into treatment and control groups and exposed to continuous noise or maintained under control conditions until the mid-hatchling stage (5 h 30 min post-fertilization at 19°C). Embryos were then collected, photographed and scored for developmental phenotypes, classified into (1) normally developed hatchlings, (2) aberrant hatchlings with visible morphological alterations, (3) pre-hatching abnormal embryos that failed to form a tailbud, and (4) un-cleavage eggs. Morphology, mortality rate and overall development were assessed using Fiji image analysis software [30]. Developmental outcomes were compared using a ratio-based approach that normalized the proportion of normally developed hatchlings in noise-exposed embryos relative to their paired controls from the same pooled clutch; adopting values near one when there were no differences, or significantly lower than one when the number of well-developed hatchlings decreased in noise treatments. After checking for normality with Shapiro–Wilk normality test, comparisons were performed using t-tests between noise-exposed embryos and paired untreated controls.

### Transcriptomic analysis

For transcriptomic experiments, total RNA from noise-treated and control-untreated embryos at 8-cell (8c, 45min pf) and early tailbud (ETB, 2h40’ pf) stages was obtained using the RNeasy Mini Kit (QIAGEN, Hilden, Germany; cat. no. 74104). Each noise-exposure experiment was replicated five times for 8c and ten times for ETB. In all cases, treated and control synchronous embryos always came from a single fertilization of a pool of eggs and sperm from various individuals to ensure biological variability (**Additional File 1: Table S1**). Samples were store at-80°C until sent in dry ice for sequencing (Novogene, Germany). Libraries were prepared following the company’s standard protocols and sequenced on the Illumina NovaSeq X Plus platform using pair-end reads of 150 bp. 8c libraries were sequenced on a single batch while ETB libraries were sequenced across two batches.

RNA-seq data was processed using the nf-core/rnaseq pipeline implemented in Nextflow, which integrates quality control, read trimming, read alignment/quantification, and gene-level summarization in a reproducible workflow [31]. The genome reference use was the one described in Plessy et al., 2024 [24] and the gene annotation was obtained with CORAL (v1.3.0; [25]) using the available long-read RNAseq data (BioProject PRJNA1347569). Global principal component analysis (PCA) was performed using the *prcomp* function in R to assess sample homogeneity and identify outlier biological replicates. Differential gene expression (DGE) analysis was performed with DESeq2 [32], filtering out genes with less than 3 samples with at least 10 mapped read counts each. An adjusted p-value of ≤ 0.05 and a |log2FoldChange| ≥ 0.6 for ETB stage and ≥ 0.2 for 8c stage were considered as the threshold for significance. Results were visualized using volcano plots and Venn diagrams generated in R with *ggplot2* [33] and *ggforce* [34], the latter using custom scripts for overlap analysis. Since ETB libraries were sequenced across two independent sequencing batches, differential expression analyses for ETB samples were first performed separately for each batch, and subsequently, by a meta-analysis approach to combine the effect sizes across batches minimizing batch-driven biases. Meta-analysis was performed in R using the *metafor* package [35], combining log2 fold changes and their associated standard errors under a random-effects model (REML estimator). Only genes detected in both batches and showing concordant directionality of expression changes were retained for downstream analyses. Lastly, heatmaps were generated in R using *pheatmap* [36] from variance-stabilized and row-scaled expression values. Top differentially expressed genes (DEGs; BH adjusted p-value < 0.05) were visualized.

### Functional annotation and validation

Functional annotation and gene ontology (GO) terms for *O. dioica* annotated genes were obtained with EggNOG-mapper [37], using the predicted proteome as input, and setting score to 40, percentage of identity, query cover, and subject cover to 20 and the e-value threshold to 0.001.

To validate and refine these functional annotations on target DEGs, their sequences were further analysed by BLASTp [38] against the NCBI non-redundant database and by InterProScan [39] for domain prediction. Genes were classified into three confidence categories based on concordance between annotation sources: (i) high-confidence annotations, supported by consistent EggNOG, BLAST and domain predictions; (ii) conserved uncharacterized proteins, lacking EggNOG annotation but showing strong BLAST similarity to homologous proteins in *Oikopleura dioica* or related species; and (iii) domain-based annotations, where functional inference relied on conserved protein domains (e.g., EGF-like motifs, glycosyltransferase domains) despite limited sequence-level annotation. Additional structural features, including signal peptides, transmembrane regions, coiled-coil domains and intrinsically disordered regions, were predicted using Phobius, SignalP and MobiDB-lite to further support functional interpretation [40–42].

### Gene Ontology enrichment analysis

GO enrichment analysis was performed implementing Over-Representation Analysis (ORA), separately for upregulated and downregulated DEGs. In all cases, the function *enricher* from the R package *clusterProfiler* [43] was implemented using as input the list of target DEGs and as reference the GO annotations obtained with EggNOG-mapper. False discovery rate (FDR; ‘BH’ parameter on *enricher*) was used as adjustment method, setting an adjusted p-value threshold of 0.05 and a minimum count of 2 for significance.

### Weighted gene co-expression network analysis

Weighted gene co-expression network analysis (WGCNA) was performed to identify modules of co-expressed genes associated with noise exposure at each developmental stage. Variance-stabilized expression data obtained with DESeq2 were used as input, and genes were filtered to retain the top 50% most variable genes. Signed co-expression networks were constructed using the R package WGCNA [44]. The soft-thresholding power was selected according to the scale-free topology criterion. Gene modules were identified by hierarchical clustering followed by dynamic tree cutting (minimum module size = 30; merge cut height = 0.25). Module eigengenes were calculated and correlated to experimental traits (noise-treated, control-untreated) using Pearson correlation, and statistical significance was assessed using asymptotic Student p-values. Functional enrichment of module-associated genes was performed using ORA as described above, using the set of expressed genes at each developmental stage as background.

## Results

### Phenotypic effects, malformation rate and viability of *O. dioica* embryo development during high intensity exposure across different noise profiles

Considering that the impact of noise on *O. dioica* had never been studied to the best of our knowledge, our first approach in this work consisted on testing a panel of different frequencies and intensities to understand in which conditions the biology of *O. dioica* could be affected, paying special attention to low frequencies as recommended by EC guidelines [45]. Given the semelparous and short life cycle of *O. dioica*, our study focused on the impact of noise pollution embryonic development because it probably constitutes the most critical and fully environmentally exposed stage that makes this species vulnerable to environmental stressors [46]. Embryos were continuously exposed from fertilization until mid-hatchling stage (5h30 min pf) to high frequencies of both pure tones (PT: 32.5 Hz, 125 Hz, 500 Hz, and 1000 Hz) and noise bands (NB: 32.5–125 Hz, 63–125 Hz, 125–500 Hz, and 500–1000 Hz), across a range of high-intensity levels (153–163 dB re 1 µPa) (**Fig. 2A**). Quantification of malformations across more than 12,000 embryos, revealed no significant differences between noise-exposed and control embryos, with developmental success values near to 1 (**Fig. 2B-C**; **Additional File 1: Table S2**). These findings, therefore, reveal that at least at the macromorphological level, early *O. dioica* embryos exhibited a high degree of resilience to noise exposure, even when exposed to the EC/NB:63–125Hz low-frequency noise range at maximum intensity, which had been predicted to represent the most widespread source of low-frequency sound in European seas according to the European Commission (EC) recommendations [45].

**Figure 2.**
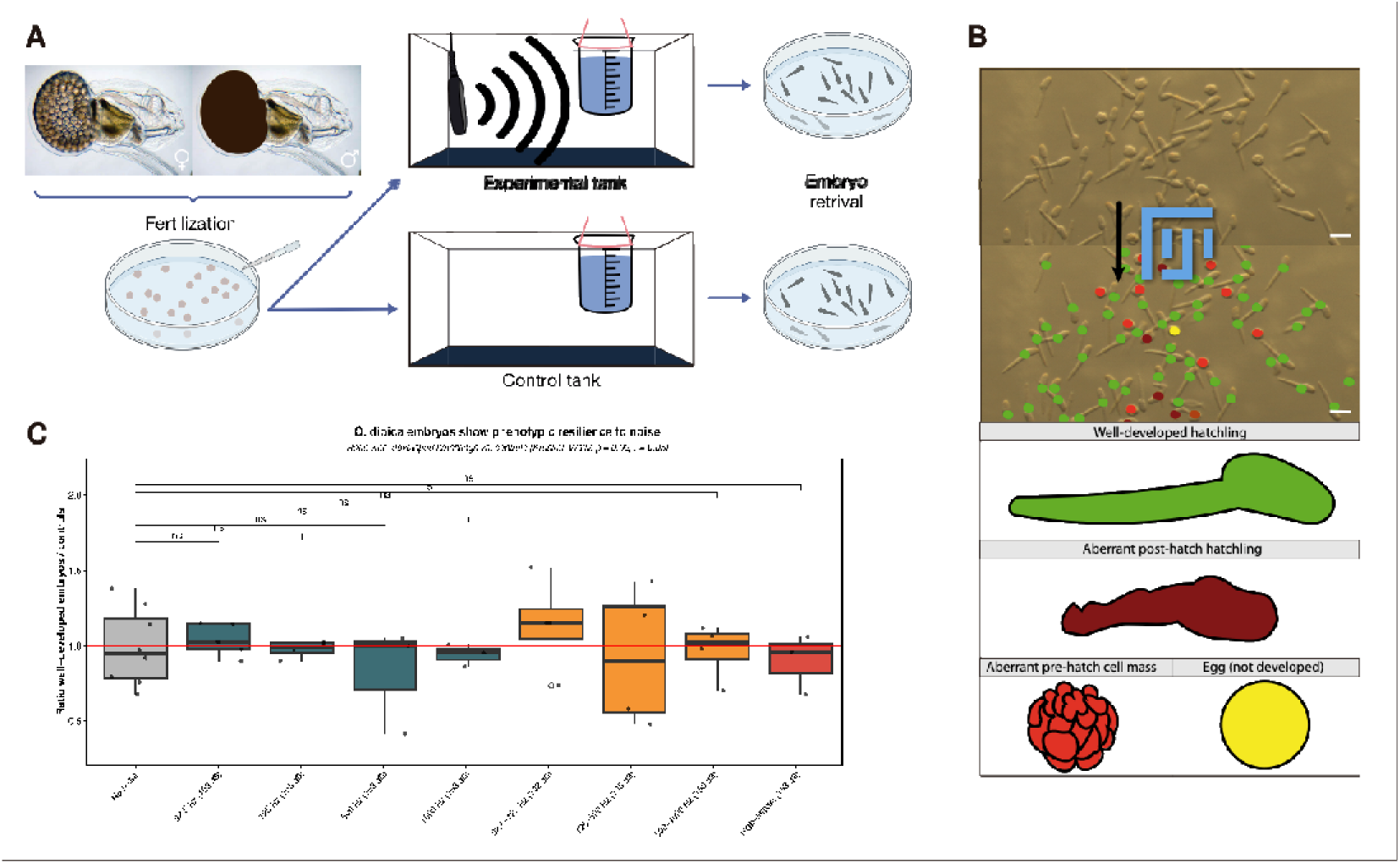
High-intensity noise exposure experiments on *O. dioica* embryos. **(a)** Protocol for noise exposure experiment. **(b)** After 5.5 h of noise exposure, embryo phenotypes could be classified in four categories: unfertilized eggs (yellow), pre-hatch arrested embryos (light red), aberrant hatchlings (dark red) and normal well-developed embryos (green). Scale bar 50 μm. **(c)** Box plot where each dot represents a replicate of the ratio of well-developed embryos of the noise-treated embryos to the controls. When the ratio is 1 there is no effect, if lower, the proportion of well-developed embryos decrease with treatment. There are at least three replicates per treatment.

### Global transcriptomic response of *O. dioica* embryos to noise exposure

To investigate the genetic molecular bases underlying the morphological resilience of *O. dioica* embryo development, we conducted a differential gene expression (DGE) analysis by a transcriptomic approach based on bulk RNA sequencing (RNA-seq) comparing the expression of embryos that had been exposed to the EC/NB:63–125Hz noise at maximum intensity (156 dB re 1 µPa) to non-exposed control embryos. Sequencing generated high-quality data across the 30 analyzed samples, with an average of 37.2 million reads per sample and a mean alignment rate of 91.9% ± 3.0% (SD) (**Additional File 1: Table S3**). Global principal component analysis (PCA) showed a clear separation of samples according to developmental stage (i.e. 8c *vs.* ETB stages), while biological replicates clustered together within stages (**Additional File 1: Fig. S2A-C**). Stage-specific PCA revealed clustering of biological replicates with moderate within-group variability, consistent with biological heterogeneity in *O. dioica* [47]. Two outliers were identified through PCA analysis. One corresponded to an 8c-stage biological replicate pair (control and treatment) that clustered separately from the remaining replicates, suggesting a slight developmental shift relative to the expected stage. A second pair, belonging to the ETB stage, followed a trajectory clearly divergent from all other replicates and was therefore excluded from downstream analyses (**Additional File 1: Fig. S2E-F**). Following removal of outliers, all experimental groups retained sufficient biological replicates with a consistent grouping by treatment within each developmental stage (**Additional File 1: Fig. S2**).

Underlying the morphological resilience of *O. dioica* embryo development to noise treatments, our DGE analyses uncovered a significant number of genes that respond with changes in their expression levels upon noise exposure (**Fig. 3; Additional File 2: Table S4**). These results revealed a rapid genetic response to noise, as a 30-minute exposure was sufficient to trigger subtle but significant transcriptional changes of 70 genes as early as the 8c stage (45’ pf). These changes were characterized by relatively small fold-change magnitudes and included 44 upregulated and 26 downregulated genes (**Table 2**; **Fig. 3A, C**). The DGE analysis on ETB stage (2h40’ pf) revealed that this initial genetic response was quickly amplified, leading to a total of 700 DEGs, the majority of which (533) were downregulated (**Table 3**; **Fig. 3B, D**).

**Figure 3.**
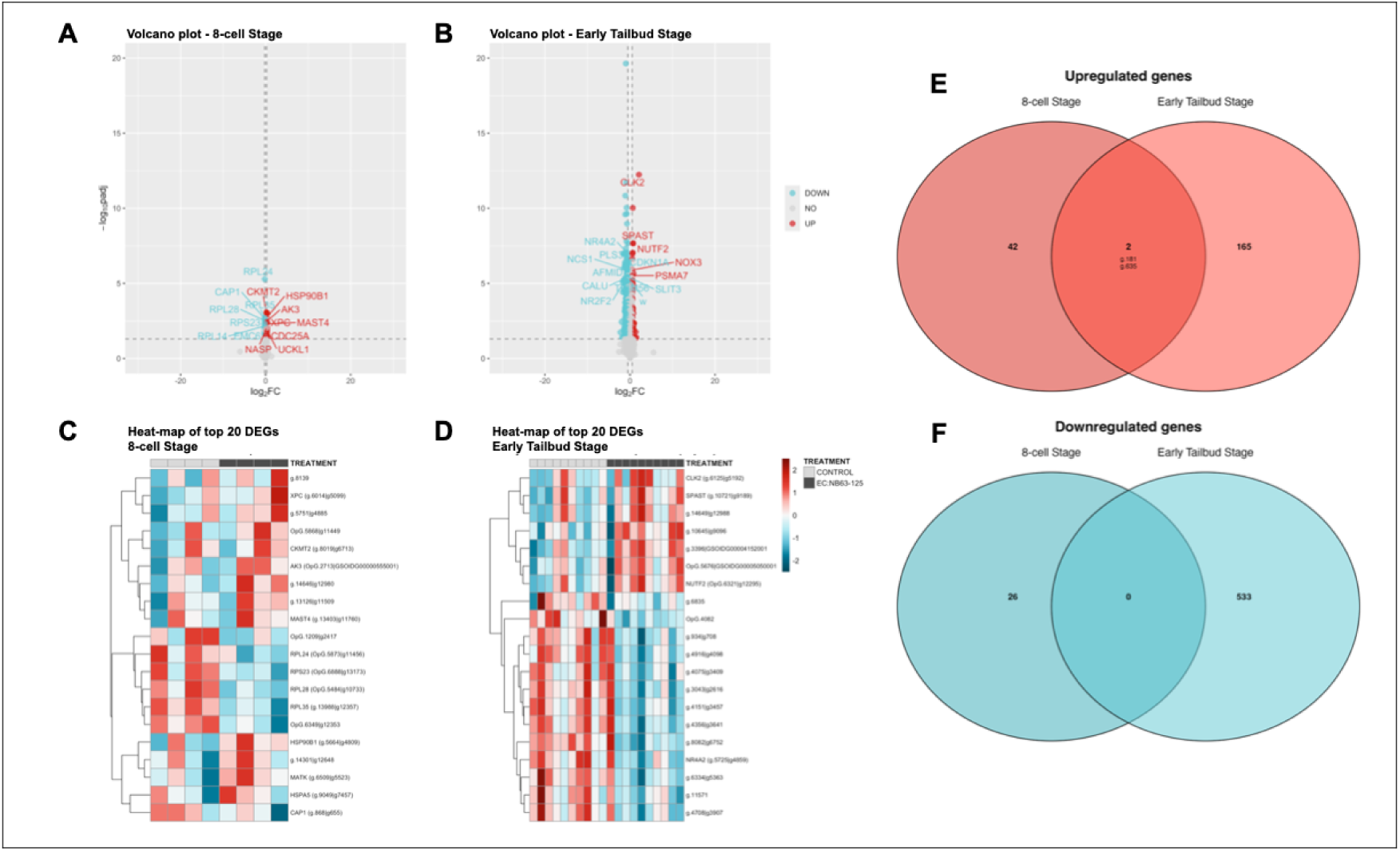
Gene Expression Dynamics in *O. dioica* Across Developmental Stages Following Noise Exposure. (a-b) Volcano plot of differentially expressed genes in 8c **(a)** and ETB **(b)** stages following noise exposure. **(c-d)** Heatmap of the 20 most significant differentially expressed genes in 8c **(c)** and ETB **(d)** stages following noise exposure. **(e-f)** Venn diagrams of up-and downregulated genes across stages, showing gene counts. Labels indicate orthologs when available; genes without orthologs retain original identifiers.

**Table 1.** Top 20 differentially expressed genes by log2FoldChange in 8c embryos. For each gene, the table shows the gene identifier, EggNOG ortholog (if available), curated functional annotation (see Methods), mean normalized expression (*baseMean*), log2 fold change (*log2FoldChange*), standard error (*lfcSE*), Wald statistic (stat), and adjusted p-value (Benjamini–Hochberg). Direction of differential expression is indicated as upregulated (UP) or downregulated (DOWN).

| Gene | EggNOG ortholog | Final annotation (category) | baseMean | log2FoldChange | lfcSE | stat | adj. p-value | Diff. expr. |
| --- | --- | --- | --- | --- | --- | --- | --- | --- |
| **g.8139** | - | RING-H2 E3 ubiquitin ligase-like (i) | 1472,23 | 0,37 | 0,08 | 4,36 | 4,39E-03 | UP |
| **OpG.3571** | CDC25A | CDC25-like (i) | 1703,87 | 0,34 | 0,08 | 4,06 | 9,25E-03 | UP |
| **g.6587** | - | SUMO-specific protease / ULP-like (i) | 1053,08 | 0,34 | 0,10 | 3,32 | 4,51E-02 | UP |
| **g.10477** | DDX6 | DDX6-like RNA helicase (i) | 1271,51 | 0,32 | 0,10 | 3,35 | 4,22E-02 | UP |
| **g.14471** | UCKL1 | UCKL1-like (i) | 4925,10 | 0,31 | 0,08 | 3,90 | 1,24E-02 | UP |
| **g.14646** | - | rhotekin-2-like (i) | 6977,53 | 0,31 | 0,07 | 4,41 | 3,84E-03 | UP |
| **OpG.5868** | - | uncharacterized secreted protein (ii) | 4427,25 | 0,30 | 0,06 | 4,97 | 8,79E-04 | UP |
| **g.14674** | - | NAV/UNC53-like (i) | 1907,62 | 0,29 | 0,07 | 3,97 | 1,06E-02 | UP |
| **g.11332** | CDC25A | CDC25-like (i) | 1817,34 | 0,29 | 0,08 | 3,78 | 1,68E-02 | UP |
| **g.7703** | - | CCCH zinc-finger RNA-binding protein (i) | 2248,16 | 0,28 | 0,07 | 4,02 | 9,69E-03 | UP |
| **OpG.991** | - | intrinsically disordered uncharacterized (ii) | 1774,79 | 0,27 | 0,07 | 3,66 | 2,26E-02 | UP |
| **g.2575** | - | mediator/RNA pol II suppressor-like (i) | 2146,28 | 0,27 | 0,08 | 3,55 | 2,89E-02 | UP |
| **g.13126** | - | bridge-like lipid transfer protein (i) | 4330,17 | 0,27 | 0,05 | 4,93 | 8,79E-04 | UP |
| **g.2819** | SLC2A5 | SLC2A5-like transporter (i) | 1625,97 | 0,26 | 0,07 | 3,57 | 2,73E-02 | UP |
| **g.5664** | HSP90B1 | HSP90B1 / GRP94-like (i) | 6840,98 | 0,26 | 0,05 | 4,71 | 1,91E-03 | UP |
| **OpG.2713** | AK3 | AK3 mitochondrial adenylate kinase (i) | 5154,27 | 0,25 | 0,06 | 4,47 | 3,37E-03 | UP |
| **g.12735** | - | coiled-coil protein (iii) | 2792,40 | 0,25 | 0,07 | 3,42 | 3,56E-02 | UP |
| **g.9704** | - | ZFAT-like zinc finger TF (i) | 1627,01 | -0,28 | 0,07 | -4,05 | 9,25E-03 | DOWN |
| **g.5176** | - | no annotation (ii) | 2928,48 | -0,26 | 0,07 | -3,78 | 1,68E-02 | DOWN |
| **OpG.6349** | - | RPS24 / eS24 (i) | 78846,48 | -0,26 | 0,06 | -4,28 | 4,93E-03 | DOWN |

Functional categories: (i) annotated genes with known or predicted function; (ii) uncharacterized or low-information genes; (iii) partial or ambiguous annotations.

**Table 2.** Top 20 differentially expressed genes by log2FoldChange in ETB embryos. Columns and annotation categories are as described in Table 2.

| Gene | EggNOG ortholog | Final annotation (category) | baseMean | log2FoldChange | lfcSE | stat | adj p-value | Diff. expr. |
| --- | --- | --- | --- | --- | --- | --- | --- | --- |
| **g.6125** | CLK2 | CLK2 (i) | 601,65 | 2,15 | 0,26 | 8,36 | 1,66E-14 | UP |
| **g.11960** | - | Nudix hydrolase-like (i) | 95,08 | 1,81 | 0,52 | 3,50 | 3,83E-03 | UP |
| **g.6554** | - | alpha-tubulin-like (partial) (iii) | 72,69 | 1,62 | 0,52 | 3,11 | 1,09E-02 | UP |
| **g.5919** | ALKBH5 | ALKBH5 (i) | 79,81 | 1,58 | 0,40 | 3,96 | 9,22E-04 | UP |
| **g.6377** | TPM3 | TPM3 (i) | 856,64 | -2,21 | 0,15 | -15,14 | 6,15E-48 | DOWN |
| **g.9150** | OTX2 | OTX/OTXa (i) | 342,22 | -2,08 | 0,20 | -10,57 | 6,15E-23 | DOWN |
| **g.2001** | - | fibrillar collagen alpha-like (i) | 9404,15 | -2,01 | 0,46 | -4,34 | 2,31E-04 | DOWN |
| **g.6571** | - | secreted extracellular protein (i) | 2318,04 | -1,85 | 0,18 | -10,54 | 7,00E-23 | DOWN |
| **OpG.408 2** | - | Oikopleura-specific gene (ii) | 182,45 | -1,82 | 0,36 | -5,10 | 9,91E-06 | DOWN |
| **g.1824** | - | secreted uncharacterized protein (ii) | 454,12 | -1,74 | 0,31 | -5,54 | 1,18E-06 | DOWN |
| **g.3898** | - | fibrillin-like ECM protein (i) | 144,23 | -1,70 | 0,19 | -9,17 | 2,28E-17 | DOWN |
| **g.1448** | PAN3 | PAN3 (i) | 605,54 | -1,70 | 0,37 | -4,59 | 9,13E-05 | DOWN |
| **g.14303** | - | Kelch/BTB-domain protein (i) | 499,51 | -1,59 | 0,21 | -7,77 | 1,32E-12 | DOWN |
| **g.1995** | - | EGF-domain protein (i) | 2406,33 | -1,58 | 0,27 | -5,87 | 2,21E-07 | DOWN |
| **g.5901** | - | Ig/Laminin-G ECM protein (i) | 416,43 | -1,58 | 0,20 | -7,79 | 1,18E-12 | DOWN |
| **g.11167** | - | myosin regulatory light chain-like (i) | 725,26 | -1,51 | 0,30 | -5,08 | 1,09E-05 | DOWN |
| **OpG.158** | - | connexin / gap junction protein (i) | 1197,97 | -1,51 | 0,34 | -4,37 | 2,14E-04 | DOWN |
| **g.3689** | KLHL24 | KLHL24 (i) | 454,70 | -1,49 | 0,37 | -4,06 | 6,58E-04 | DOWN |
| **g.9327** | - | tubulin alpha chain (i) | 391,06 | -1,48 | 0,16 | -8,96 | 1,17E-16 | DOWN |
| **g.12926** | ESRP2 | ESRP2 (i) | 1035,10 | -1,45 | 0,21 | -6,91 | 4,70E-10 | DOWN |

Functional categories: (i) annotated genes with known or predicted function; (ii) uncharacterized or low-information genes; (iii) partial or ambiguous annotations.

**Table 3.** Common upregulated genes in both 8c and ETB embryos. Columns and annotation categories are as described in Table 2.

| Gene | EGGNOG ortholog | Final annotation (category) | baseMean | log2Fold Change | lfcSE | stat | adj p-value | Stage |
| --- | --- | --- | --- | --- | --- | --- | --- | --- |
| **g.635** | - | weak similarity to motile sperm uncharacterized protein (iii) | 6236,32 | 0,25 | 0,07 | 3,67 | 2,21E-02 | 8c |
|  |  |  | 1439,77 | 0,63 | 0,24 | 2,61 | 3,35E-02 | ETB |
| **g.181** | - | P-loop NTPase-like (ii) | 4575,87 | 0,24 | 0,07 | 3,31 | 4,51E-02 | 8c |
|  |  |  | 990,19 | 0,64 | 0,19 | 3,32 | 6,32E-03 | ETB |

Functional categories: (i) annotated genes with known or predicted function; (ii) uncharacterized or low-information genes; (iii) partial or ambiguous annotations.

To explore the biological functions associated with the observed genetic response, we first performed Gene Ontology (GO) enrichment analyses by ORA on the up-and downregulated DEGs identified at each developmental stage. Then, to capture coordinated transcriptional patterns beyond individual gene-level changes, we applied weighted gene co-expression network analysis (WGCNA), enabling the identification of modules of co-expressed genes and their associated biological functions. Thus, while ORA identified significantly over-represented pathways based on differentially expressed genes, WGCNA captured coordinated expression patterns at the network level, providing a complementary systems-level perspective (**Fig. 4**; **Table 4**).

**Figure 4.**
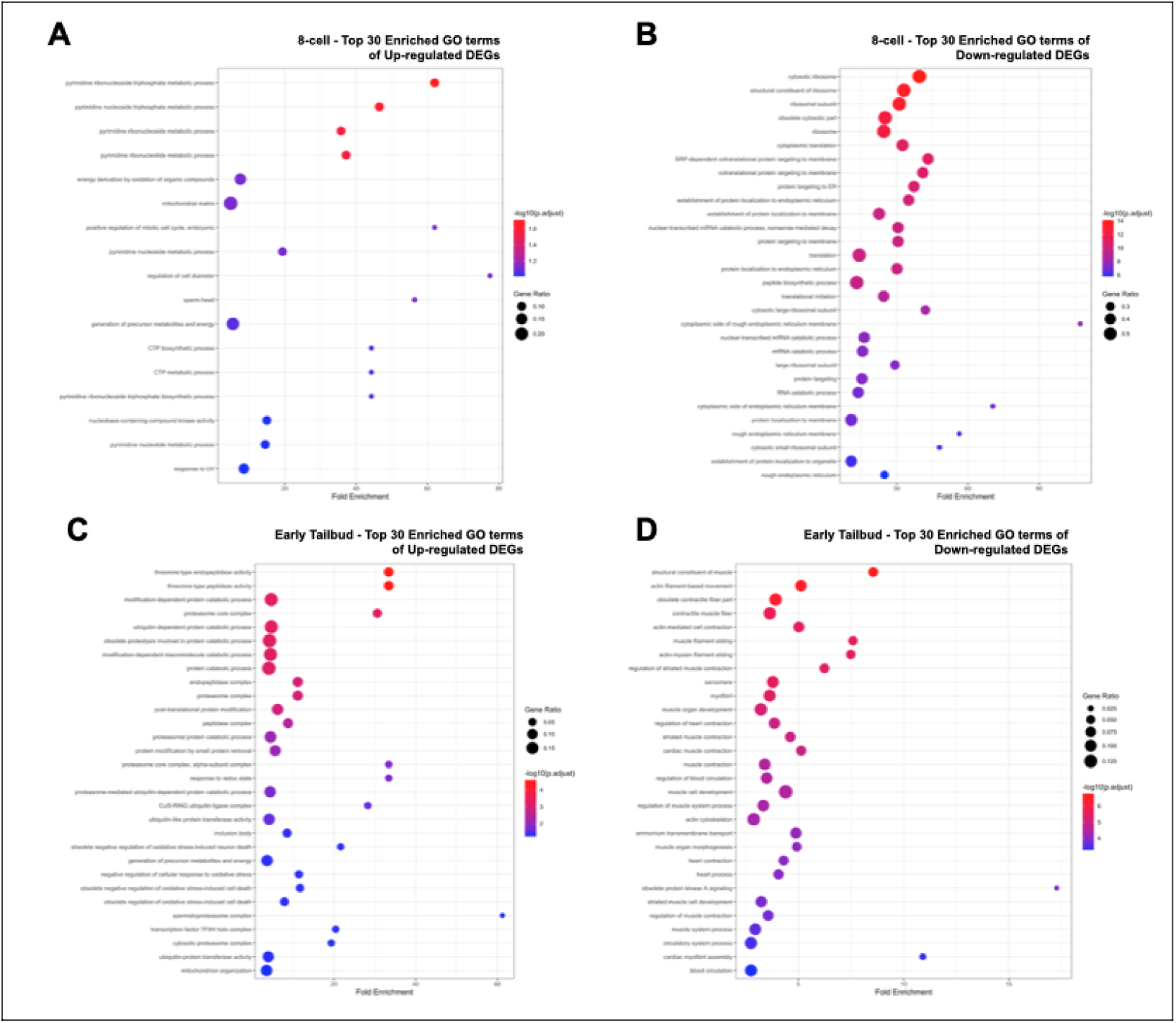
**Enrichment analyses in *O. dioica* Across Developmental Stages Following Noise Exposure**. **(a-b)** Enriched GO terms in 8c stage following noise exposure. **(c-d)** Enriched GO terms in ETB stage following noise exposure.

**Table 4.**
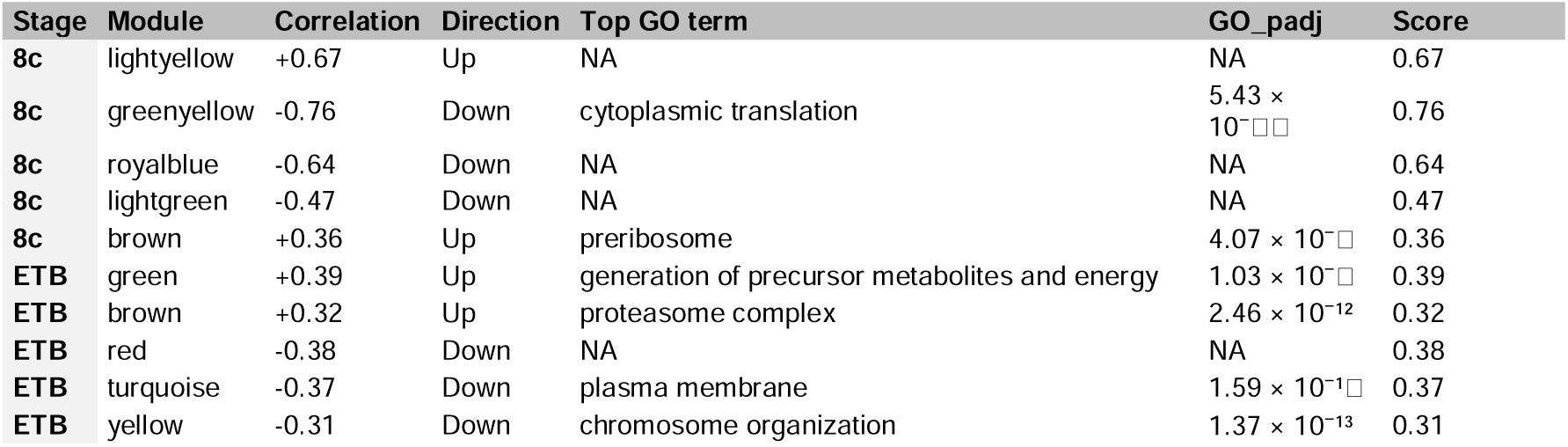
Integrated summary of the most biologically relevant noise-associated modules identified by WGCNA. . Modules are ranked according to their combined score, integrating module–trait correlation strength and GO enrichment significance. For each module, developmental stage, correlation with noise, direction of association, top enriched GO term, and adjusted p-value are shown.

At 8c stage, GO enrichment analysis revealed that many of enriched terms among upregulated DEGs were associated with CTP and pyrimidine ribonucleoside metabolic processes (**Fig. 4A; Additional File 2: Table S5**). Upregulation of CTP and pyrimidine ribonucleoside has been described to be not only important for increased RNA synthesis in situations of high transcription demand or cell proliferation, but also critical for lipid biogenesis associated to cellular membrane dynamics [48]. Indeed, CTP has been described to be central for the maintenance of lipid membrane homeostasis and repair, being activated by stress-induced elastic membrane curvatures [49]. This result, therefore, plausibly suggested that CTP activity upregulation could be related to noise-induced perturbation of membrane homeostasis or membrane-associated cellular dynamics of embryos as soon as 30 minutes of exposure at the 8c stage. In this context, we also found as enriched terms among upregulated DEGs cell diameter, sperm-head proteins related to membrane protrusions, as well as embryonic cell cycle, mitochondrial and energetic metabolism. On the other hand, amongst the downregulated DEGs, we found many enriched GO terms that related to protein targeting to membranes and protein localization in rough endoplasmic reticulum, which again suggested that noise could be affecting cellular membrane dynamics. We also observed amongst downregulated DEGs enrichment of GOs terms related to ribosome subunit assembly and protein translation, suggesting that noise could also be affecting internal membranes, impacting therefore other fundamental processes such as translation and transcription (**Fig. 4B**).

At the ETB stage, the number of enriched GO terms increased approximately five-fold, reaching a total of 234 terms. Among terms enriched among upregulated DEGs, many reflected the activation of protein degradation by proteostasis and ubiquitin–proteasome pathways. (**Fig. 4C; Additional File 2: Table S5**). Activation of protein degradation pathway could be indicative that noise could also promote protein misfolding or proteostatic imbalance, which therefore could drive the fate of those misfolded proteins to be actively degraded. Enrichment analysis also suggested that noise-induced protein alterations were coupled with oxidative stress responses, post-translational protein regulation, and metabolic/mitochondrial remodelling associated with cellular stress adaptation. Among downregulated DEGs, most enriched terms were associated with transcriptional regulation, extracellular matrix (ECM) and cytoskeleton organization, and cell fate and differentiation, particularly muscle and cardiac development (**Fig. 4D**). These enriched terms could be reflecting how noise could ultimately affect fundamental cellular and developmental processes.

WGCNA further supported the coordinated nature of these transcriptional responses to noise exposure (**Table 4; Additional File 2: Table S6-7**). In both developmental stages, modules positively correlated with noise exposure were associated with metabolic, mitochondrial, and cellular stress-response functions, whereas negatively correlated modules were enriched in processes related to translation, membrane organization, signalling, and developmental cellular functions, suggesting a broad reallocation of cellular resources in response to acoustic stress. However, stage-specific differences were also evident. At the 8c stage, the strongest negatively correlated modules were primarily enriched in translation-, ribosome-, and protein synthesis-related functions, indicating an early repression of translational activity shortly after noise exposure. In contrast, positively correlated modules at the ETB stage were more strongly associated with mitochondrial, metabolic, and intracellular regulatory processes, while negatively correlated modules were enriched in membrane-and signalling-related functions, pointing to a more functionally specialized cellular response during later embryonic development.

Overall, these results indicated that noise enabled a quick first genetic response as soon as only 30 minutes after exposure already at 8c stage, which was already rapidly amplified after 2h of exposure by specially affecting cellular membrane dynamics and protein degradation as two potential direct noise-specific consequences that could be influencing early embryo development.

### Top-listed genes related to noise in *O. dioica* embryos

At the gene level, several of the top DEGs (**Tables 1, 2**) further reinforced the biological interpretations emerging from the GO enrichment analyses, particularly those related to membrane homeostasis, protein degradation, and developmental cellular organization. At the 8c stage, one of the most informative upregulated genes was a bridge-like lipid transfer protein 2 (BLTP2-like; g.13126), a family of proteins associated with lipid transfer between intracellular membranes and membrane-contact site organization. Another informative upregulated gene showed sequence similarity to Uridine-cytidine kinases (UCKL-1 like; g.14471), which has been described to play important roles in the pyrimidine salvage pathway [50]. The upregulation of these two genes provided additional support to the enrichment of CTP and pyrimidine-related metabolic processes, compatible with the hypothesis that noise exposure might rapidly induce cellular responses associated with membrane remodelling and lipid homeostasis. Supporting this interpretation, we also identified the upregulation of a rhotekin-2-like gene (g.14646), involved in Rho-mediated cortical actin and membrane dynamics, and the NAV/UNC53 (g.14674), which has been associated to the regulation of cytoskeleton dynamics connected to endocytotic active cells [51]. Together, these genes suggested that noise exposure could rapidly affect cortical and membrane-associated cellular dynamics, including the cytoskeleton, during the earliest stages of embryogenesis.

Several genes also supported the activation of proteostasis, protein degradation and protein quality-control pathways. Among them, a RING-H2 E3 ubiquitin ligase-like gene (g.8139) represented one of the strongest direct indicators of ubiquitin-mediated protein degradation, while the upregulation of a SUMO-specific protease/SENP-like gene (g.6587) further pointed to the activation of stress-responsive post-translational regulatory systems. In addition, embryos exposed to noise showed increased expression of an HSP90B1/GRP94-like chaperone (g.5664), an endoplasmic reticulum-associated molecular chaperone involved in protein folding and unfolded protein responses. Altogether, these genes strongly supported the enrichment of proteostasis-and ubiquitin–proteasome-related GO terms, suggesting that noise exposure increases the demand for protein quality-control and cellular homeostasis pathways.

The 8-cell-stage response also included several genes associated with RNA metabolism and cell-cycle regulation, consistent with the enrichment of metabolic and translational regulatory functions. These included a *DDX6-like* RNA helicase (g.10477), involved in maternal mRNA turnover and translational repression and two *CDC25*-like phosphatases (OpG.3571 and g.11332), which has been associated to the regulation of embryonic cell-cycle progression. The coordinated differential expression of multiple RPL genes further supported the early impact of noise exposure on translational machinery and ribosome-related functions at the 8c stage (**Additional File 2: Table S4**). Together, the upregulation of these genes suggested that noise exposure rapidly triggers subtle but coordinated regulatory adjustments affecting RNA metabolism, translation, and cleavage-stage cellular dynamics.

This regulatory signature became more pronounced at the ETB stage, where genes such as the CDC Like Kinase 2 (*CLK2*, g.6125), involved in RNA splicing regulation, the RNA demethylase *ALKBH5* (g.5919), associated with epitranscriptomic control of RNA stability and translation, and a Nudix hydrolase-like gene (g.11960), potentially linked to nucleotide and RNA quality control under cellular stress pointed to an expanded post-transcriptional and homeostatic response and supports the broader enrichment of stress adaptation, proteostasis, and metabolic regulatory pathways identified in the functional analyses. However, the transcriptional response became substantially broader and was characterized predominantly by downregulation of genes associated with ECM organization, cytoskeletal architecture, and developmental regulation. Among the most strongly downregulated genes were several ECM-related proteins, including fibrillar collagen alpha-1-like (g.2001), EGF-containing fibrillin-like extracellular matrix protein (g.3898), and Ig/Laminin-G domain-containing extracellular matrix protein (g.5901), together with additional secreted extracellular proteins (g.6571 and g.1824). This coordinated downregulation strongly supported the GO enrichment analyses indicating perturbation of ECM organization and tissue morphogenesis.

Similarly, several cytoskeleton-related genes were also downregulated at ETB stage, including tropomyosin 3 (*TPM3*; g.6377), tubulin alpha chain (g.9327), and a myosin regulatory light chain-like gene (g.11167), suggesting alterations in cellular architecture and morphogenetic dynamics. The downregulation of PAN3 (g.1448), which has been described as a regulatory subunit of the deadenylation complex associated to functions of mRNA stabilization suggest that noise might also interfere with the machinery of mRNA post-transcriptional regulation. In parallel, developmental regulatory genes such as *OTXa* (g.9150), a conserved developmental transcription factor that is already early expressed during gastrulation [52], and *ESRP2* (g.12926), an epithelial splicing regulator involved in epithelial differentiation and morphogenesis, were also strongly downregulated. These results suggested that prolonged exposure to noise not only affects cellular homeostasis pathways but may ultimately interfere with developmental regulatory programs and tissue organization during embryogenesis.

Comparison of common DEGs between 8c and ETB stages revealed two shared genes, despite these ones were not in the top-list of each stage (**Fig. 3E**; **Table 3**). The first gene, g.635, appeared annotated as an unknown gene, but interestingly, BLAST analysis reveals that this gene had orthologs exclusively within the tunicate subphylum, but not in other clades, suggesting that it could be an evolutionary innovation of this clade. InterProScan analysis revealed the presence of an MSP (motile sperm protein) domain, which has been described as the signature of a family of proteins than function in cell motility independent of actin and tubulin by changing cytoskeleton organization, and promoting cell membrane protrusions that drive ameboid-like cell movements [53]. The second gene, g.181, also appeared annotated as an unknown gene, and only having clear orthologs in *O. dioica* genomes. InterProScan analysis revealed the presence of a P-loop containing nucleoside triphosphate hydrolase domain, which is characteristic of proteins than can hydrolase nucleosides for energy obtention that could be used for remodelling protein complexes, RNA/DNA processing, membrane trafficking, cytoskeletal dynamics [54].

Overall, the combined GO enrichment and gene-level analyses supported a model in which noise exposure first induces rapid cellular homeostatic responses involving membrane remodelling, protein degradation and proteostasis, RNA regulation, and metabolic adaptation, which later become associated with broader perturbations affecting ECM organization, cytoskeletal dynamics, and developmental regulatory pathways during embryonic morphogenesis.

## Discussion (UNDER CONSTRUCTION)

### Morphological resilience to noise during embryo development of *O. dioica*

Understanding how marine invertebrates respond to environmental stressors, including the growing impact of marine noise pollution, is essential for assessing the vulnerability of marine ecosystems and, ultimately, ocean health. This is particularly relevant for zooplanktonic organisms, which occupy central positions in marine trophic webs and contribute fundamentally to nutrient cycling and carbon fluxes in the oceans. Among them, appendicularian jelly tunicates, including the cosmopolitan species *O. dioica,* represent especially important components of pelagic ecosystems [55]. Through their highly efficient filter-feeding activity and the continuous production of discarded mucus houses and fecal pellets, appendicularians contribute substantially to marine snow formation and carbon sequestration, playing a key role in the biological carbon pump [26].

Beyond their ecological importance, appendicularians also represent an informative model for investigating environmental perturbations because of their rapid life cycle and semelparous reproductive strategy. In *O. dioica*, individuals reproduce only once and die immediately after releasing gametes, meaning that reproductive success fully depends on a single developmental cycle. Consequently, disturbances affecting embryogenesis cannot be compensated by subsequent reproductive events, potentially making early embryos particularly vulnerable to environmental stressors. Since embryogenesis occurs fully exposed in the surrounding marine environment, understanding how embryos respond to anthropogenic noise becomes especially relevant for evaluating the biological consequences of acoustic pollution in marine ecosystems [46].

To our knowledge, this work represents the first study investigating the impact of noise exposure on *O. dioica* embryogenesis. Despite the absence of significant increases in overt developmental abnormalities, noise exposure elicited clear and progressive transcriptional responses, indicating that embryos rapidly sense and process acoustic perturbation at the molecular level. This apparent resilience of *O. dioica* embryos contrasts with observations reported in other marine organisms, where anthropogenic noise has been associated with reduced embryonic survival, impaired hatching success, developmental delays, and morphological abnormalities. For example, decreased embryo viability has been documented in sea hares (*Stylocheilus striatus*) and cuttlefish (*Sepia officinalis*) exposed to anthropogenic noise, while impulsive acoustic exposure has been linked to developmental abnormalities and delayed growth in scallop larvae (*Pecten novaezelandiae*) and Norway lobster (*Nephrops norvegicus*) larvae [56–59].

The coexistence of morphological robustness and widespread transcriptional remodeling suggests that the absence of visible developmental defects does not necessarily reflect absence of biological impact. Instead, embryos may maintain developmental integrity through compensatory cellular responses that reallocate metabolic and regulatory resources toward stress adaptation and homeostasis [60, 61]. Similar phenotype–molecular mismatches have increasingly been recognized in environmental stress biology, where transcriptomic responses often provide earlier and more sensitive indicators of perturbation than organism-level traits alone [62, 63]. In *O. dioica*, such resilience may additionally be facilitated by the species’ rapid r-selected life cycle and remarkable genomic plasticity, including extensive genome reduction, operon organization, and exceptionally high rates of genome rearrangement [24].

### Genetic response of *O. dioica* early embryos to noise

The embryonic stages analyzed here precede the formation of differentiated mechanosensory structures potentially associated with sound perception. Consequently, the observed transcriptional responses are unlikely to reflect sensory-mediated pathways comparable to those operating in later developmental stages or adults. Instead, our results suggest that acoustic exposure may directly perturb cellular homeostasis, triggering measurable molecular responses within only 30 minutes of exposure.

Among the biological processes affected by noise exposure, two emerging categories appeared particularly informative because they may reflect direct cellular consequences of acoustic stress: membrane homeostasis and mechanical organization, and proteostasis-related pathways associated with protein folding, degradation, and quality control.

-*Noise and membrane homeostasis*

One of the strongest signatures emerging from our analyses involved processes associated with membrane homeostasis and cellular mechanical organization. Upregulation of CTP-and pyrimidine ribonucleoside-related pathways, together with enrichment of membrane-associated processes, may indicate rapid modulation of membrane homeostasis following acoustic exposure. Beyond its role in nucleotide metabolism and RNA synthesis, CTP is central to phospholipid biosynthesis and membrane maintenance, participating in lipid remodeling and membrane repair processes activated under conditions affecting membrane deformation or membrane tension [49, 48].

This interpretation is consistent with the growing recognition that cellular membranes and associated cytoskeletal systems act as primary interfaces for mechanosensation and mechanotransduction [64, 65]. Mechanical and vibrational stimulation can alter membrane curvature, lipid organization, cytoskeletal tension, and membrane-associated signaling pathways, thereby influencing downstream cellular responses [66]. Consistent with this mechanobiological framework, ETB embryos displayed coordinated regulation of extracellular matrix and contractility-associated genes, including collagen-, fibrillin-, laminin-, and cytoskeleton-related components, whereas 8c embryos showed modulation of genes potentially associated with membrane organization and Rho-mediated cortical tension signaling. Similar structural and ECM-associated responses have also been reported in other marine organisms exposed to anthropogenic noise [17, 18], suggesting that cellular mechanical remodeling may represent a conserved component of acoustic stress responses across phylogenetically distant taxa.

Importantly, this mechanobiological framework may help explain aspects of the transcriptional response that appear less compatible with classical chemical-stress signatures. Many pollutants, particularly toxicants and xenobiotics, commonly induce responses dominated by oxidative stress, detoxification pathways, xenobiotic metabolism, and inflammatory signaling. In contrast, mechanically induced stress responses more frequently affect extracellular matrix organization, membrane dynamics, cytoskeletal architecture, and force-transduction pathways [64, 65].

-*Noise and proteostasis*

A second major response involved activation of protein degradation, proteostasis and cellular quality-control pathways. Noise exposure induced regulation of molecular chaperones, ubiquitin-associated regulators, SUMO-related pathways, and multiple RNA-regulatory factors, indicating coordinated activation of systems involved in protein folding, RNA processing, and cellular maintenance [67]. Particularly notable was the repression of translation-and ribosome-associated pathways, especially during early exposure stages. Both GO and WGCNA analyses identified coordinated downregulation of ribosome biogenesis, ribosomal assembly, and translational processes, further supported by differential expression of multiple ribosomal protein genes during the 8-cell response. Suppression of translation is a common compensatory adaptive strategy during environmental stress, allowing cells to reduce energetic expenditure while redirecting resources toward homeostasis, repair, and stress-response functions [68, 69]. Under isolated laboratory conditions, these compensatory mechanisms may be sufficient to preserve morphological stability despite substantial transcriptional reorganization. However, under natural multi-stressor environments combining noise with warming, pollutants, microplastics, or marine biotoxins, such buffering mechanisms may become insufficient, potentially amplifying physiological and developmental disruption during embryogenesis

## Conclusions

Early embryos of *O. dioica* display a rapid and progressively amplified transcriptional response to underwater noise exposure despite showing strong morphological resilience, highlighting the value of this species as an emerging model for studying acoustic stress responses in marine zooplankton.

Differential gene expression analyses suggest that noise exposure induces coordinated cellular stress responses compatible with mechanobiological perturbation, particularly affecting membrane homeostasis, proteostasis, and protein degradation pathways. Together, these responses are consistent with compensatory stress-adaptation mechanisms aimed at preserving cellular integrity and developmental stability during embryogenesis.

## Supporting information

Addtional_files_1_2

## List of abbreviations

8c: 8-cell
DEG: Differentially Expressed Gene
DGE: Differential Gene Expression
EC: European Commission
ECM: Extracellular Matrix
ETB: Early tailbud
FDR: False Discovery Rate
GO: Gene Ontology
NB: Noise Band
ORA: Over-Representation Analysis
PCA: Principal Component Analysis
PT: Pure Tone
WGCNA: Weighted gene co-expression network analysis

## Declarations

### Ethics approval and consent to participate

Not applicable

### Consent for publication

Not applicable

### Availability of data and materials

The RNA-seq datasets generated during the current study are available in the Sequence Read Archive (SRA) database under accession numbers SRR38812248 – SRR38812277, BioProject PRJNA1469892. All data generated or analysed during this study are included in this published article and its supplementary information files. Custom scripts used for data analyses are available from the corresponding author on reasonable request.

### Authors’ contributions

E.R.Q. led the wet and computational experiments, performed the analyses, created the figures and wrote the manuscript. N.P.T.A. contributed to obtaining *O. dioica* material, data curation, computational interpretations and manuscript writing and editing. I.N.P. performed the acoustic calibration of the experimental setup. S.N., V.C., G.B., M.B., E.P., G.S., I.N.P. and E.R.Q. contributed to the experiment design. E.R.Q., S.C., I.G., L.M., R.P., F.R., G.Z., M.C., R.M.A.P. and C.C. conceived the study within the Deuteronoise consortium. R.M.A.P. and C.C. acquired funding, supervised the research, and contributed to the interpretation of the results and manuscript writing. All authors read and approved the final manuscript.

## Acknowledgements

The authors thank the ‘Centres Científics i Tecnològics de la UB’ for sea water supply, Novogene for their sequencing services and, especially, Sebastian Artime Paoletti for running the Oikopleura facility at the University of Barcelona, the other members of the EvoDevo Genomics UB lab for helpful discussions, and Josep F. Abril Ferrando for his support on our computational server at the UB.

