## Supplementary material for "Developmental genetic response of the zooplanktonic tunicate *Oikopleura dioica* to marine noise pollution": Addtional_files_1_2: Additional_File_1.pdf

**Table S1. Statistical analysis of developmental success across acoustic treatments.** For each condition (*sound file*), the table reports the number of replicates (*n*), results of the Shapiro–Wilk normality test (*Shapiro-Wilk p-value*), raw p-values and Benjamini–Hochberg adjusted p-values (*p-value, adjusted p-value*), and statistical significance. Comparisons were performed using t-tests between noise-exposed embryos and paired untreated controls. ns: not significant.

| Sound condition | n | Shapiro–Wilk p-value | p-value | adjusted p-value | Significance |
|---|---|---|---|---|---|
| **No noise** | 8 | 0.6206063 | 0.9180656 | 0.9180656 | ns (p=0.918) |
| **PT32.5** | 5 | 0.4503151 | 0.4596060 | 0.7264949 | ns (p=0.46) |
| **PT125** | 4 | 0.3259552 | 0.4843299 | 0.7264949 | ns (p=0.484) |
| **PT500** | 3 | 0.1395621 | 0.4723632 | 0.7264949 | ns (p=0.472) |
| **PT1000** | 3 | 0.7712217 | 0.3054329 | 0.7264949 | ns (p=0.305) |
| **NB32.5-125** | 4 | 0.6758482 | 0.4489828 | 0.7264949 | ns (p=0.449) |
| **EN:NB63-125** | 3 | 0.46294251 | 0.4702299 | 0.7264949 | ns (p=0.47) |
| **NB125-500** | 4 | 0.34381018 | 0.7641446 | 0.8596627 | ns (p=0.764) |
| **NB500-1000** | 4 | 0.33083832 | 0.7390800 | 0.8596627 | ns (p=0.739) |

**Table S2. Sample metadata and RNA quality.** For each sample, the table shows the number of animals pooled (*females* + *males*), estimated number of eggs, total RNA yield (*ng*), and RNA integrity (*RIN*) values, reflecting the quality of RNA used for downstream analyses.

| Sample Name | Animals pooled | Nº of eggs (estimate) | RNA yield (ng) | RNA integrity (RIN) |
|---|---|---|---|---|
| **HINTc_ETB_R1** | 3♀+6♂ | 225 | 430 | 9,8 |
| **HINTn_ETB_R1** | | 225 | 390 | 9,8 |
| **HINTc_ETB_R2** | 4♀+3♂ | 300 | 100 | 9,9 |
| **HINTn_ETB_R2** | | 300 | 430 | 9,9 |
| **HINTc_ETB_R3** | 3♀+3♂ | 300 | 290 | 9,8 |
| **HINTn_ETB_R3** | | 300 | 630 | 9,2 |
| **HINTc_ETB_R4** | 3♀+3♂ | 300 | 280 | 9,7 |
| **HINTn_ETB_R4** | | 300 | 620 | 9,9 |
| **HINTc_ETB_R5** | 3♀+3♂ | 300 | 510 | 9,8 |
| **HINTn_ETB_R5** | | 300 | 500 | 9,9 |
| **HINTc_ETB_R6** | 9♀+5♂ | 675 | 320 | 9,9 |
| **HINTn_ETB_R6** | | 675 | 380 | 9,8 |
| **HINTc_ETB_R7** | 9♀+4♂ | 675 | 1430 | 9,9 |
| **HINTn_ETB_R7** | | 675 | 1645,5 | 9,9 |
| **HINTc_ETB_R8** | 8♀+4♂ | 600 | 470 | 9,9 |
| **HINTn_ETB_R8** | | 600 | 666 | 9,9 |
| **HINTc_ETB_R9** | 8♀+4♂ | 600 | 830 | 9,9 |
| **HINTn_ETB_R9** | | 600 | 2500,5 | 9,9 |
| **HINTc_ETB_R10** | 8♀+5♂ | 600 | 440 | 9,9 |
| **HINTn_ETB_R10** | | 600 | 1047 | 9,9 |
| **HINTc_8c_R1** | 8♀+4♂ | 600 | 814 | 9,9 |
| **HINTn_8c_R1** | | 600 | 960 | 9,9 |
| **HINTc_8c_R2** | 9♀+8♂ | 675 | 1430 | 9,9 |
| **HINTn_8c_R2** | | 675 | 1480 | 9,9 |
| **HINTc_8c_R3** | 8♀+6♂ | 600 | 220 | 9,9 |
| **HINTn_8c_R3** | | 600 | 370 | 9,9 |
| **HINTc_8c_R4** | 8♀+4♂ | 600 | 130 | 9,9 |
| **HINTn_8c_R4** | | 600 | 160 | 9,9 |
| **HINTc_8c_R5** | 9♀+5♂ | 675 | 1880 | 9,9 |
| **HINTn_8c_R5** | | 675 | 1190 | 9,9 |

**Table S3. Sequencing depth and alignment quality metrics for all RNA-seq samples.** For each sample, the table shows total reads generated, overall alignment rate (*Aligned*), uniquely aligned reads (*Uniq aligned*), average mapped read length (*Avg. mapped len*), number of annotated splice junctions (*Annotated splices*), and rates and lengths of mismatches, deletions, and insertions (*Mismatch rate, Del rate, Del len, Ins rate, Ins len*) observed during alignment.

| Sample Name | Total reads | Aligned | Uniq aligned | Avg. mapped len | Annotated splices | Mismatch rate | Del rate | Del len | Ins rate | Ins len |
|---|---|---|---|---|---|---|---|---|---|---|
| **HINTc_ETB_R1** | 31.1M | 91.5% | 89.9% | 293.6bp | 29.3M | 0.7% | 0.0% | 2.1bp | 0.0% | 1.8bp |
| **HINTc_ETB_R2** | 38.3M | 89.9% | 88.2% | 293.5bp | 34.4M | 0.7% | 0.0% | 2.1bp | 0.0% | 1.7bp |
| **HINTc_ETB_R3** | 45.7M | 90.9% | 89.2% | 293.9bp | 42.3M | 0.7% | 0.0% | 2.1bp | 0.0% | 1.8bp |
| **HINTc_ETB_R4** | 37.2M | 93.4% | 91.8% | 293.5bp | 35.6M | 0.7% | 0.0% | 2.1bp | 0.0% | 1.8bp |
| **HINTc_ETB_R5** | 42.3M | 90.9% | 89.3% | 293.9bp | 41.1M | 0.7% | 0.0% | 2.1bp | 0.0% | 1.8bp |
| **HINTc_ETB_R6** | 32.8M | 83.5% | 81.9% | 289.7bp | 26.3M | 0.7% | 0.0% | 2.1bp | 0.0% | 1.7bp |
| **HINTc_ETB_R7** | 29.8M | 87.8% | 85.9% | 290.9bp | 25.1M | 0.8% | 0.0% | 2.2bp | 0.0% | 1.8bp |
| **HINTc_ETB_R8** | 35.1M | 96.1% | 94.2% | 295.4bp | 33.3M | 0.8% | 0.0% | 2.1bp | 0.0% | 1.8bp |
| **HINTc_ETB_R9** | 39.2M | 96.0% | 94.3% | 293.3bp | 38.5M | 0.7% | 0.0% | 2.1bp | 0.0% | 1.8bp |
| **HINTc_ETB_R10** | 38.9M | 95.8% | 94.0% | 294.1bp | 38.4M | 0.7% | 0.0% | 2.1bp | 0.0% | 1.8bp |
| **HINTn_ETB_R1** | 35.5M | 91.4% | 89.9% | 292.5bp | 34.4M | 0.6% | 0.0% | 2.1bp | 0.0% | 1.8bp |
| **HINTn_ETB_R2** | 45.8M | 91.7% | 90.1% | 294.1bp | 43.1M | 0.7% | 0.0% | 2.1bp | 0.0% | 1.8bp |
| **HINTn_ETB_R3** | 31.4M | 89.2% | 87.7% | 292.8bp | 28.9M | 0.7% | 0.0% | 2.1bp | 0.0% | 1.8bp |
| **HINTn_ETB_R4** | 33.0M | 92.9% | 91.5% | 294.6bp | 32.4M | 0.6% | 0.0% | 2.1bp | 0.0% | 1.9bp |
| **HINTn_ETB_R5** | 33.4M | 89.5% | 88.0% | 294.2bp | 32.7M | 0.7% | 0.0% | 2.1bp | 0.0% | 1.8bp |
| **HINTn_ETB_R6** | 33.0M | 93.9% | 92.2% | 294.1bp | 30.6M | 0.7% | 0.0% | 2.1bp | 0.0% | 1.7bp |
| **HINTn_ETB_R7** | 36.8M | 94.7% | 92.9% | 293.5bp | 35.4M | 0.7% | 0.0% | 2.2bp | 0.0% | 1.8bp |
| **HINTn_ETB_R8** | 47.3M | 93.8% | 92.1% | 293.0bp | 44.2M | 0.7% | 0.0% | 2.1bp | 0.0% | 1.8bp |
| **HINTn_ETB_R9** | 36.9M | 92.9% | 91.3% | 293.1bp | 35.2M | 0.7% | 0.0% | 2.1bp | 0.0% | 1.8bp |
| **HINTn_ETB_R10** | 37.5M | 87.8% | 86.2% | 291.6bp | 34.0M | 0.7% | 0.0% | 2.1bp | 0.0% | 1.8bp |
| **HINTc_8c_R1** | 35.9M | 92.2% | 90.9% | 292.5bp | 36.8M | 0.6% | 0.0% | 2.1bp | 0.0% | 1.8bp |
| **HINTc_8c_R2** | 36.0M | 93.0% | 91.6% | 292.8bp | 36.9M | 0.6% | 0.0% | 2.1bp | 0.0% | 1.7bp |
| **HINTc_8c_R3** | 38.4M | 92.8% | 91.4% | 292.9bp | 39.5M | 0.6% | 0.0% | 2.1bp | 0.0% | 1.8bp |
| **HINTc_8c_R4** | 35.6M | 92.4% | 91.1% | 293.3bp | 36.2M | 0.6% | 0.0% | 2.1bp | 0.0% | 1.8bp |
| **HINTc_8c_R5** | 36.4M | 90.1% | 88.9% | 292.8bp | 38.3M | 0.6% | 0.0% | 2.0bp | 0.0% | 1.9bp |
| **HINTn_8c_R1** | 35.3M | 92.9% | 91.7% | 292.0bp | 36.2M | 0.6% | 0.0% | 2.1bp | 0.0% | 1.8bp |
| **HINTn_8c_R2** | 38.2M | 94.3% | 93.0% | 292.8bp | 39.9M | 0.6% | 0.0% | 2.1bp | 0.0% | 1.8bp |
| **HINTn_8c_R3** | 45.4M | 90.2% | 88.9% | 292.5bp | 44.9M | 0.6% | 0.0% | 2.1bp | 0.0% | 1.8bp |
| **HINTn_8c_R4** | 36.5M | 90.6% | 89.3% | 292.9bp | 36.2M | 0.6% | 0.0% | 2.2bp | 0.0% | 1.8bp |
| **HINTn_8c_R5** | 31.9M | 91.2% | 89.9% | 292.8bp | 35.3M | 0.6% | 0.0% | 2.0bp | 0.0% | 1.8bp |

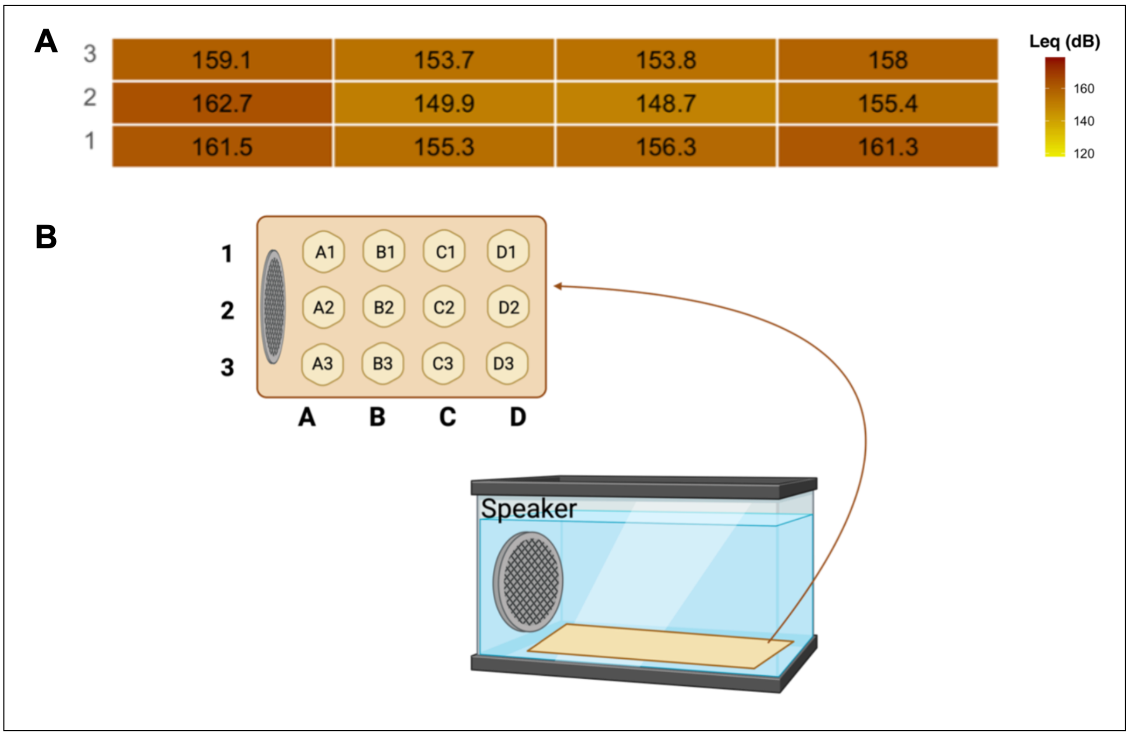


**Figure S1: Tank Acoustic Calibration. (a)** Heatmap of a pink noise sound file with energy peaks at 63 and 125 Hz, levels (Leq, 30min) inside the glass tank across 12 measurements (sites 1-3 x A-D). The equivalent noise level (Leq), necessary for the subsequent evaluations, was calculated by processing the measurement files using specific software (SpectraPlus-SC), considering a time interval of 30 seconds in which the recording was as steady as possible for a more significant result. During noise treatments the speaker was positioned in 2A. **(b)** Scheme of the placement of the measurements made in the tank, diving the tank into a grid of 3x4, both inside the treatment and control. Using this knowledge, we took measurements on the specific site where the animal was fixed in the tank while exposed to noise for the different playbacks (C2).

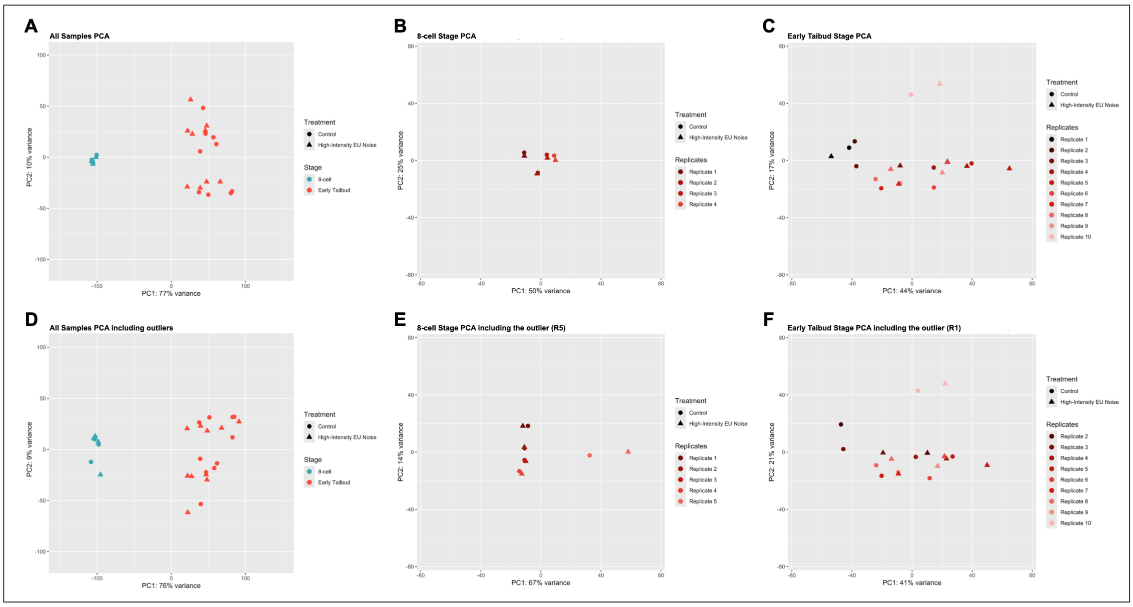


**Figure S2. Principal component analysis (PCA) of transcriptomic profiles. (a)** PCA including all developmental stages. **(b–c)** Stage-specific PCAs for samples exposed to EC:NB63–125 at the 8-cell and Early Tailbud stages, respectively. **(d–f)** PCA plots before removal of outlier samples, including all developmental stages **(d)**, 8-cell embryos **(e)**, and Early Tailbud embryos **(f)**. Outlier samples corresponded to R5 in the 8-cell dataset and R1 in the Early Tailbud dataset.
